# Quantitative interactome proteomics identifies proteostasis network for GABA_A_ receptors

**DOI:** 10.1101/2022.03.08.483512

**Authors:** Ya-Juan Wang, Xiao-Jing Di, Ting-Wei Mu

## Abstract

Gamma-aminobutyric acid type A (GABA_A_) receptors, the primary inhibitory neurotransmitter-gated ion channels in the mammalian central nervous system, inhibit neuronal firing to preserve balanced neuronal activity. Maintenance of GABA_A_ receptor protein homeostasis (proteostasis) in the cell utilizing its interacting proteins is essential for the function of GABA_A_ receptors. However, how the proteostasis network orchestrates GABA_A_ receptor biogenesis in the endoplasmic reticulum (ER) is not well understood. To address this question systematically, we employed a proteomics-based approach to identify the interactomes of GABA_A_ receptors by carrying out a quantitative immunoprecipitation-tandem mass spectrometry (IP-MS/MS) analysis utilizing stable isotope labeling by amino acids in cell culture (SILAC). To enhance the coverage and reliability of the identified proteins, we performed comparative proteomics by using both wild type α1 subunit and a misfolding-prone α1 subunit carrying the A322D variant as the bait proteins. The wild type α1 interactome contains 125 proteins, the α1(A322D) interactome contains 105 proteins, and 54 proteins overlap within two interactomes. Bioinformatics analysis identified potential GABA_A_ receptor proteostasis network components, including chaperones, folding enzymes, trafficking factors, and degradation factors. Further, their potential involvement is modelled in the cellular folding, degradation and trafficking pathways for GABA_A_ receptors. In addition, we verified endogenous interactions between α1 subunit and their selected interactors by carrying out co-immunoprecipitation assay in mouse brain homogenates. This study paves the way for understanding the molecular mechanisms as well as fine-tuning of GABA_A_ receptor proteostasis to ameliorate related neurological diseases such as epilepsy.

## Introduction

Normal organismal physiology depends on the maintenance of protein homeostasis (proteostasis) in each cellular compartment (1-4), which dictates a delicate balance between protein synthesis, folding, assembly, trafficking, and degradation while minimizing misfolding and aggregation (5-7). For one specific client protein, its interaction with a network of proteins, especially its proteostasis network components, in the crowded cellular environment is critical to maintain its proteostasis. However, how the proteostasis network orchestrates the biogenesis of multi-subunit multi-span ion channel proteins is poorly understood. The current limited knowledge about such protein quality control (QC) machinery is gained from the study of various classes of membrane proteins, including cystic fibrosis transmembrane conductance regulator (CFTR) (8), T cell receptors (9), sodium channels (10), potassium channels (11), and nicotinic acetylcholine receptors (12). We have been using γ-aminobutyric acid type A (GABA_A_) receptors as an important membrane protein substrate to clarify its biogenesis pathway (13), which is currently understudied.

GABA_A_ receptors are the primary inhibitory neurotransmitter-gated ion channels in mammalian central nervous systems (14) and provide most of the inhibitory tone to balance the tendency of excitatory neural circuits to induce hyperexcitability, thus maintaining the excitatory-inhibitory balance (15). Loss of their function is a prominent cause of genetic epilepsies, and recent advances in genetics have identified an growing number of epilepsy-associated variants in over ten *GABR* genes, including over 150 variants in those encoding major synaptic subunits (α1, β2, β3 and γ2 subunits) (16-20). GABA_A_ receptors belong to the Cys-loop superfamily of ligand-gated ion channels. They are pentameric, sharing common structural characteristics with other Cys-loop receptor members (21). Each subunit has a large extracellular (or the endoplasmic reticulum (ER) luminal) N-terminus, four transmembrane helices (TM1-TM4, with TM2 domain lining the interior of the pore), and a short extracellular (or the ER luminal) C-terminus (**Figure 1A**) (22,23). GABA binding to GABA_A_ receptors in the extracellular domain induces conformational changes, opens the ion pore to conduct chloride, hyperpolarizes the plasma membrane, and inhibits neuronal firing in mature neurons.

**Figure 1.**
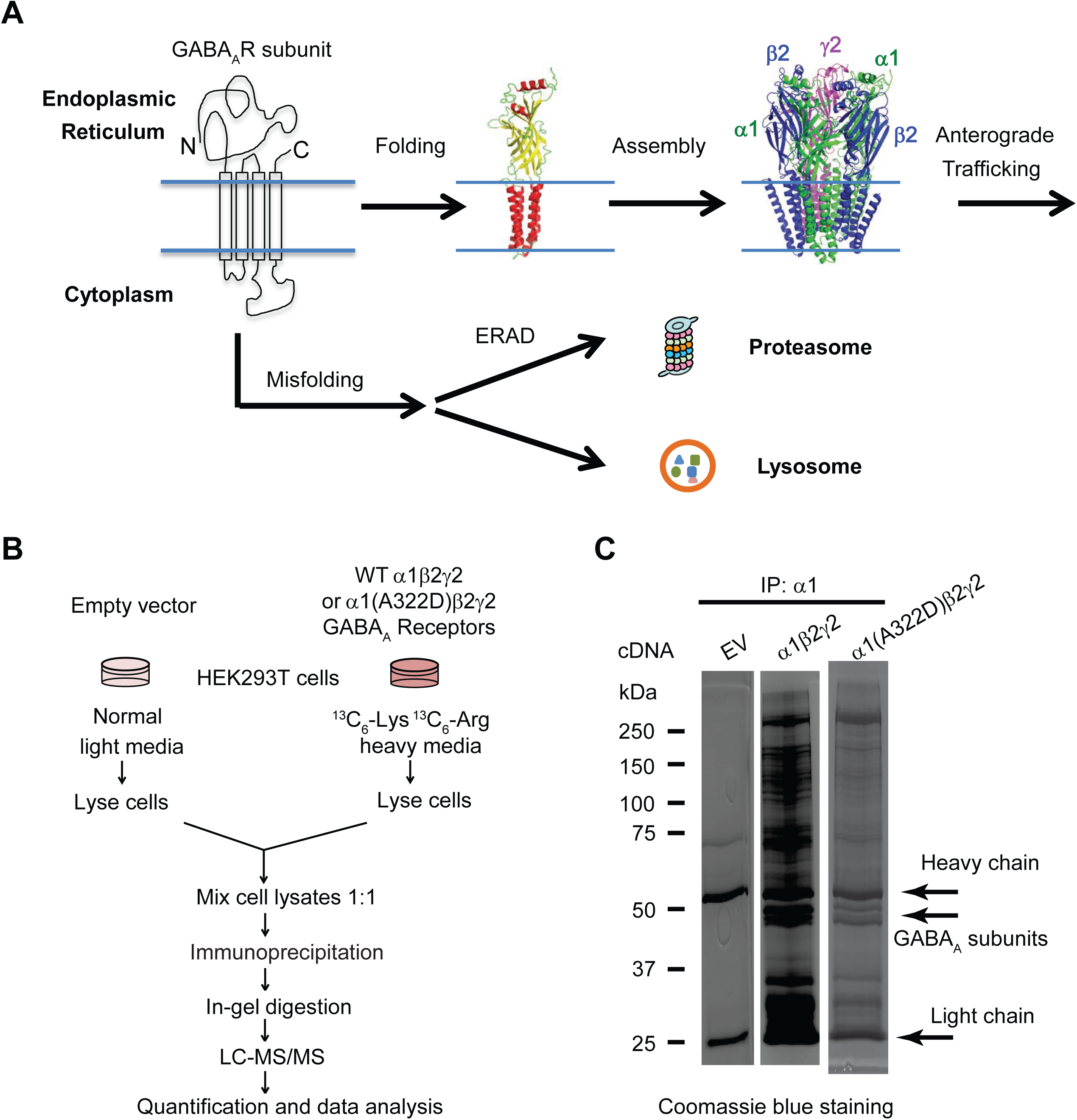
Quantitative immunoprecipitation-tandem mass spectrometry (IP-MS/MS) analysis to identify GABA_A_ receptor interactomes. (**A**) GABA_A_ receptor biogenesis pathways. Individual subunits of GABA_A_ receptors fold in the endoplasmic reticulum (ER). Properly folded subunits assemble into a heteropentamer in the ER for subsequent trafficking to the plasma membrane. Unassembled and misfolded subunits are subjected to the ER-associated degradation (ERAD) pathway by the proteasome or the lysosome-related degradation. (**B**) Outline of a comparative SILAC-based quantitative proteomics approach to identify GABA_A_ receptor interactomes in HEK293T cells. The α1(A322D) variant leads to its extensive misfolding and ERAD. (**C**) Visualization of immunoisolated wild-type (WT) and α1(A322D)β2γ2 GABA_A_ receptor complexes by SDS-PAGE and Coomassie staining. EV, empty vector; IP, immunoprecipitation.

To function properly, GABA_A_ receptors need to fold into their native structures and assemble correctly to form a pentamer on the ER membrane and traffic efficiently through the Golgi en route to the plasma membrane (**Figure 1A**). Misfolded GABA_A_ receptors are recognized by the cellular protein quality control machinery. ER-associated degradation (ERAD) is one major cellular pathway to target misfolded GABA_A_ receptors to the cytosolic proteasome for degradation (7,24). Another potential degradation pathway is to target the aggregation-prone GABA_A_ receptors to the lysosome through autophagy, ER-phagy, or ER-to-lysosome-associated degradation (ERLAD) (25,26). Maintenance of a delicate balance between GABA_A_ receptor folding, trafficking, and degradation utilizing its interacting proteins is critical for its function. However, the GABA_A_ receptor interactome, especially the proteostasis network that orchestrates GABA_A_ receptor biogenesis in the ER, has not been studied systematically in the literature despite recent advances about the trafficking of GABA_A_ receptors beyond the ER (27-31). Here, we used quantitative immunoprecipitation-tandem mass spectrometry (IP-MS/MS) analysis utilizing stable isotope labeling by amino acids in cell culture (SILAC) in HEK293T cells to identify the interactomes for both wild type and a misfolding-prone GABA_A_ receptor. Endogenous interactions between selected interactors and GABA_A_ receptors were verified in mouse brain homogenates. Furthermore, bioinformatics analysis enabled us to assemble a proteostasis network model for the cellular folding, assembly, degradation, and trafficking pathways of GABA_A_ receptors.

## Results

### Identification of interactomes for wild type and misfolding-prone GABA_A_ receptors using comparative SILAC-based proteomics

We employed a proteomics-based approach to identify the interactomes of GABA_A_ receptors in HEK293T cells by carrying out a quantitative IP-MS/MS analysis utilizing SILAC (**Figure 1B**) (32). To enhance the coverage and reliability of the identified proteins, we performed comparative proteomics by using both wild type (WT) α1 subunit and a well-characterized misfolding-prone α1 subunit carrying the A322D variant as the bait proteins (33). The A322D variant introduces an extra negative charge in the third transmembrane (TM3) helix of the α1 subunit, causing its substantial misfolding and excessive degradation by ERAD (33,34). HEK293T cells stably expressing either WT α1β2γ2 or α1(A322D)β2γ2 GABA_A_ receptors were labeled with heavy media, whereas HEK293T cells that were transfected with empty vector (EV) plasmids were cultured in normal light media. The same amount of light and heavy cell lysates was mixed. The α1 or α1(A322D) complexes were immunoprecipitated using a monoclonal antibody against the N-terminus region of the α1 subunit before being subjected to SDS-PAGE and in-gel digestion and tandem MS analysis. Coomassie blue-stained gels showed numerous clearly visible bands, including those at ∼50 kDa, which is the molecular weight for GABA_A_ receptor subunits, indicating efficient co-immunoprecipitations of the GABA_A_ receptors (**Figure 1C**).

The α1 subunit interactomes were identified using the SILAC ratio with arbitrary yet strict criteria to remove potential false positives. To be included as an interactor, it must (1) have a SILAC ratio of WT α1/EV or α1(A322D)/EV to be at least 1.30; (2) have a *p* < 0.05; and (3) have a Benjamini and Hochberg correction (35) of false discovery rate (FDR) of no more than 0.10. The top right green area in **Figure 2A** contains high-confidence interactors for GABA_A_ receptors. As a result of the stringent criteria, the WT α1 interactome contains 125 proteins, the α1(A322D) subunit interactome contains 105 proteins, and 54 proteins overlap within two interactomes (**Figure 2B)**. These 176 interactors for GABA_A_ receptors are used for the following bioinformatics analysis (see Supplemental **Table S1** for protein list).

**Figure 2.**
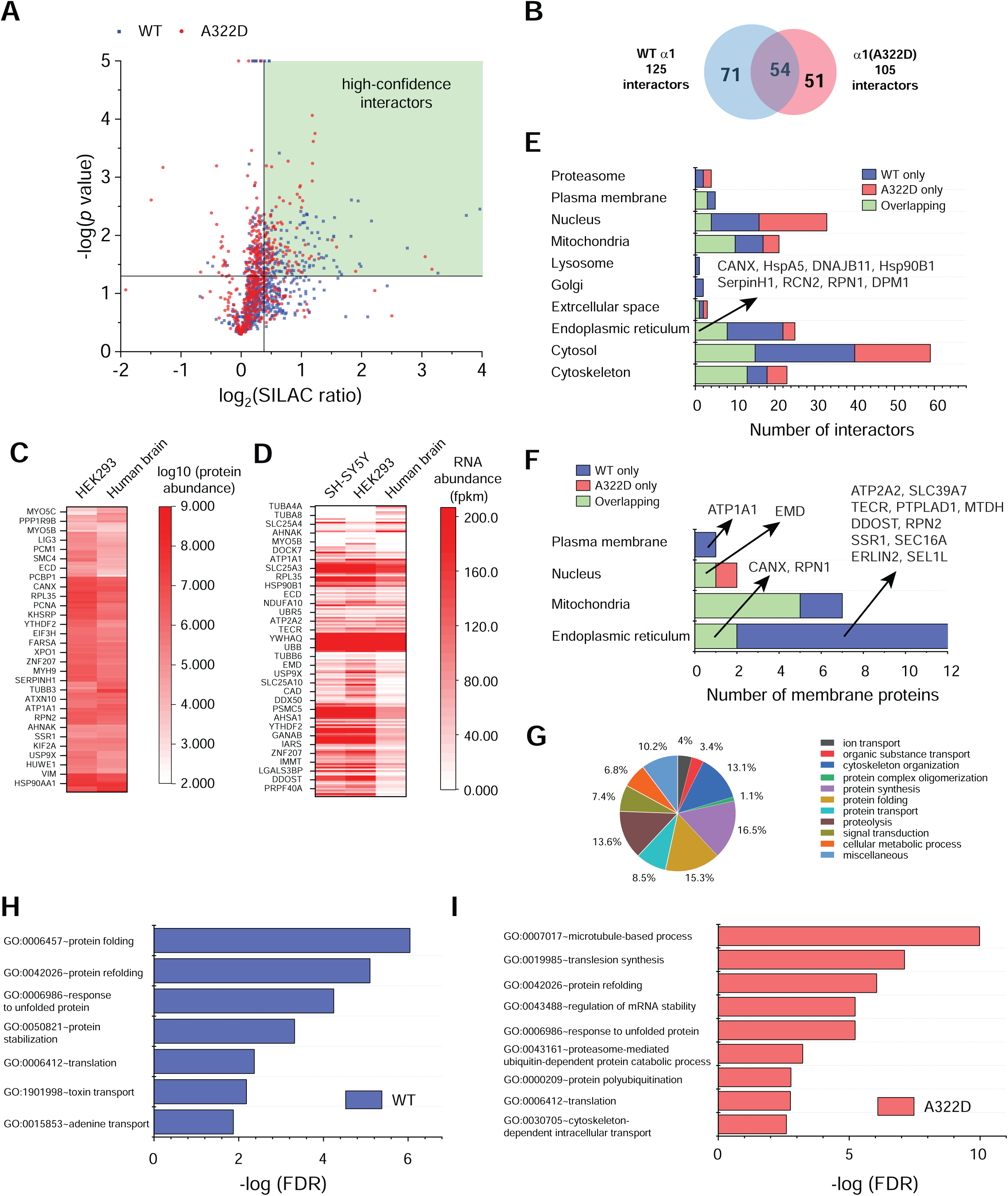
Bioinformatics analysis of GABA_A_ receptor interactomes. (**A**) A 2D plot showing the relationship between *p* value and SILAC ratio for proteins that were identified from tandem MS analysis. Proteins that are in the WT samples are colored in blue; those in α1(A322D) variant samples, in red. The vertical black line represents a SILAC ratio of 1.30, whereas the horizontal black line represents a *p* value of 0.05. (**B**) A Venn diagram showing the protein numbers and overlap of the GABA_A_ receptor interactomes. (**C**) A heat map showing the protein abundance for GABA_A_ receptor interactors in HEK293 cells and human brain tissues. (**D**) A heat map showing the RNA abundance for GABA_A_ receptor interactors in HEK293 cells, SH-SY5Y cells, and human brain tissues. Fpkm, fragments per kilobase of exon per million mapped fragments. (**E**) Analysis of cellular components for GABA_A_ receptor interactomes. (**F**) Analysis of cellular components for GABA_A_ receptor interactors that are integral membrane proteins. (**G**) A pie chart showing the biological processes of the pooled 176 GABA_A_ receptor interactors. Enriched biological processes for WT GABA_A_ receptor interactome (**H**) and α1(A322D)-containing GABA_A_ receptor interactome (**I**) according to DAVID analysis. FDR, false discovery rate.

### Gene Ontology analysis of GABA_A_ receptor interactomes

Since GABA_A_ receptors inhibit neuronal firing in the mammalian central nervous system, we first compared the expression abundance of their interactors between HEK293 cells and the nervous system. ProteomicsDB (https://www.proteomicsdb.org/) provides a mass spectrometry-based navigation of the human proteome from tissues, cell lines and body fluids, enabling the comprehensive mapping of the protein abundance of GABA_A_ receptor interactors (36). Hierarchical clustering analysis showed that these interactors have comparable protein levels between HEK293 cells and human brain tissues (**Figure 2C** and **Table S2**). For example, many molecular chaperones, such as Hsp90s (*Hsp90AA1, Hsp90AB1*, and *Hsp90B1*), Hsp70s (*HspA5, HspA8*), Hsp40s (*DNAJA1, DNAJA2, DNAJB11*), calnexin (*CANX*), and Hsp47 (*SerpinH1*), are abundantly expressed in both systems. In addition, tissue-based map of the human proteome based on quantitative transcriptomics enabled us to compare RNA levels of GABA_A_ receptor interactors between HEK293 cells, a human neuronal SH-SY5Y cell line, and human brain tissues (37,38). Hierarchical clustering analysis showed that many interactors, including molecular chaperones and ubiquitin-dependent degradation factors, such as *UBA1, UBR5, UBE3C, SEL1L*, and *VCP*, have similar RNA expression patterns between these systems (**Figure 2D** and **Table S2**). These results are consistent with the report that most proteins are well conserved among human tissues (38).

To annotate cellular component for GABA_A_ receptor interactors, we used Database for Annotation, Visualization and Integrated Discovery (DAVID) to carry out gene ontology (GO) analysis (39,40). Since many proteins reside in more than one cellular location, we only choose one primary subcellular location for them manually. To aid such an assignment, we also integrate subcellular location information from Uniprot Database (https://www.uniprot.org/) and GeneCards: The Human Gene Database (https://www.genecards.org/). GABA_A_ receptor interactors are distributed in various cellular locations, including the nucleus (33 interactors), ER (25 interactors), Golgi (2 interactors), proteasome (4 interactors), lysosome (1 interactor), mitochondria (21 interactors), cytoskeleton (23 interactors), cytosol (59 interactors), plasma membrane (5 interactors), and extracellular space (3 interactors) since biogenesis and function of GABA_A_ receptors require their interactions with a network of proteins throughout the cell (**Figure 2E** and **Table S3**). Due to the essential role of the ER in protein quality control, 25 interactors (14 for WT only, 3 for α1(A322D) variant only, and 8 for both) are located to this organelle. Eight overlapping interactors in the ER include molecular chaperones (*CANX, HspA5, DNAJB11, Hsp90B1, SerpinH1*), factors involved in *N*-linked glycosylation (*RPN1, DPM1*), and a Ca^2+^-binding protein (*RCN2*). *RCN2* encodes reticulocalbin 2 in the ER lumen; it is abundantly expressed in the brain and contains EF-hand Ca^2+^-binding motifs. Function of RCN2 is largely unknown, although gene expression of *RCN2* is upregulated in patients with idiopathic absence epilepsies (41). Given the critical role of GABA_A_ receptors in the pathophysiology of epilepsy, it will be of great interest to determine how RCN2 regulates GABA_A_ receptor biogenesis and contributes to epilepsy phenotypes in the future.

Furthermore, since GABA_A_ receptors are transmembrane proteins, we extracted their interactors that are integral membrane proteins, including 13 in the ER, 7 in the mitochondria (*SLC25A3, SLC25A4, SLC25A5, SLC25A6, SLC25A10, IMMT, TIMM50*), 2 in the nucleus (*EMD, LBR*), and 1 in the plasma membrane (*ATP1A1*) (**Figure 2F** and **Table S3**). *ATP1A1* encodes α1 subunit of sodium/potassium-transporting ATPase in the plasma membrane. Variants in *ATP1A1* cause hypomagnesemia, seizures, and mental retardation 2, an autosomal dominant disease characterized by generalized seizures in infancy and significant intellectual disability (42). The 13 membrane interactors in the ER include molecular chaperones and folding enzymes (*CANX, DDOST, RPN1, RPN2*), trafficking factors (*SSR1, SEC16A*), ERAD factors (*ERLIN2, SEL1L*), transporters (*ATP2A2, SLC39A7*), enzymes involved in the lipid metabolism (*TECR, PTPLAD1*), and other (*MTDH*). Since each subunit of GABA_A_ receptors has four transmembrane helices, these ER membrane interactors, such as *ATP2A2, SLC39A7, TECR, PTPLAD1*, and *MTDH*, have the potential to form intra-membrane interactions with GABA_A_ receptors to regulate their biogenesis in the ER, which needs to be explored in the future.

Moreover, we used DAVID, Uniprot, and GeneCards to annotate biological process for GABA_A_ receptor interactors. The 176 interactors were annotated to the following functional categories: ion transport (4%), organic substance transport (3.4%), cytoskeleton organization (13.1%), protein complex oligomerization (1.1%), protein synthesis (16.5%), protein folding (15.3%), protein transport (8.5%), proteolysis (13.6%), signal transduction (7.4%), cellular metabolic process (6.8%), and miscellaneous (10.2%) (**Figure 2G** and **Table S4**). Furthermore, we used DAVID to determine enriched biological process for GABA_A_ receptor interactors. Both WT interactors and α1(A322D) interactors form five functional clusters (**Table S5**). Top enriched biological processes from those clusters were plotted in **Figure 2H** and **Figure 2I** according to false discovery rate (FDR). Protein refolding (GO: 0042026), response to unfolded protein (GO: 0006986), and translation (GO: 0006412) are common enriched biological processes for both WT and α1(A322D) interactomes. Since the A322D variant causes extensive protein misfolding and ERAD of the α1 subunit, the α1(A322D) interactome enriches proteasome-mediated ubiquitin-dependent protein catabolic process (GO:0043161) and protein polyubiquitination (GO:0000209).

### Mapping of the proteostasis network for GABA_A_ receptors

We focused on identifying the proteostasis network for GABA_A_ receptors, which regulates their folding, assembly, trafficking and degradation. Based on GO analysis and literature knowledge, we assigned the interactors to proteostasis network categories, including protein folding (GO:0006457), proteolysis (GO:0006508), and protein transport (GO:0015031) (**Table S6**). The proteostasis network components account for 37.4% of the total interactors, indicating their critical role in regulating GABA_A_ receptor function. The number of interactors that belong to the proteostasis network was plotted for WT interactome, α1(A322D) interactome, and overlapping interactome (**Figure 3A**). In addition, we visualized the proteostasis network for WT and α1(A322D) variant-containing receptors in a 2D plot showing their SILAC ratios (**Figure 3B** and **Table S1**).

**Figure 3.**
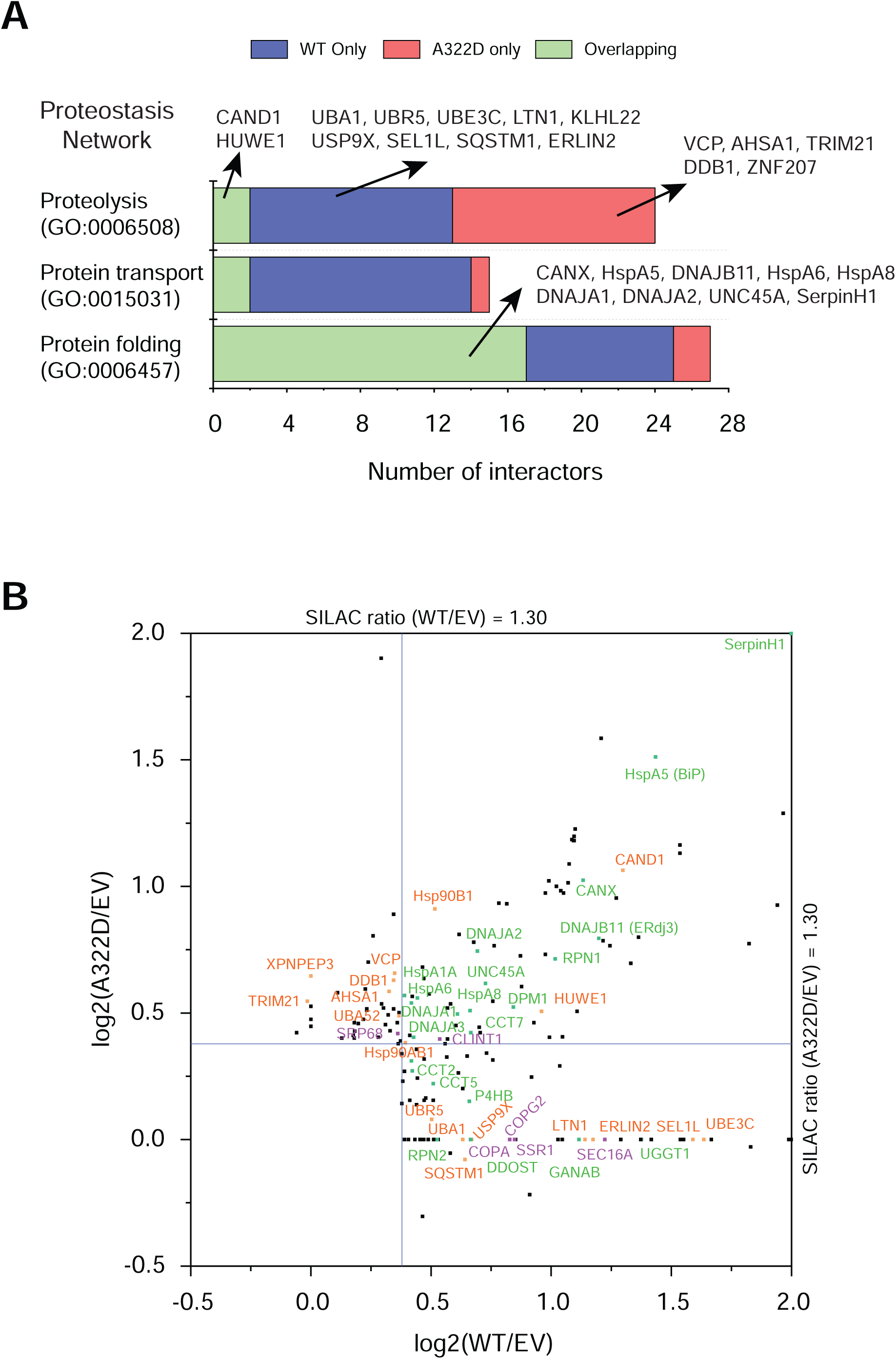
Proteome profiling of proteostasis network for WT and misfolding-prone α1(A322D)-containing GABA_A_ receptors. (**A**) Gene ontology analysis of the proteostasis network for GABA_A_ receptors interactomes. Number of interactors is plotted against proteolysis, protein transport, and protein folding. (**B**) A 2D plot showing SILAC ratios of the identified interactors. The x and y axes show log_2_(SILAC ratio of WT α1/EV) and log_2_(SILAC ratio of α1(A322D)/EV). Blue lines represent a cutoff SILAC ratio of 1.30. Based on literature knowledge, interactors that are expected to play a role in the folding and assembly of GABA_A_ receptors are colored in green; those in their degradation, in orange; and those in their transport, in purple. EV, empty vector.

Judging from the literature knowledge, interactors that are expected to regulate the folding and assembly of α1 subunits are indicated in green, those for protein transport in purple, and those for proteolysis in orange (**Figure 3B**). Major chaperone networks and folding enzymes were identified from GABA_A_ receptor interactomes. These include Hsp70s and their co-chaperone Hsp40s in the ER (HspA5 and DNAJB11) and in the cytosol (HspA6, HspA8, HspA1A, DNAJA1, DNAJA2, and DNAJA3), Hsp90s and their co-chaperones in the ER (Hsp90B1) and in the cytosol (Hsp90AB1, AHSA1, and UNC45A), Hsp60 subunits (CCT2, CCT5, and CCT7), and a protein disulfide isomerase (P4HB). Moreover, Since GABA_A_ receptors have several *N*-linked glycosylation sites, proteins that are involved in the *N*-linked glycoprotein maturation in the ER were identified, including glycosylation enzymes (DDOST, RPN1, RPN2, DPM1, GANAB, and UGGT1) and a lectin chaperone (CANX). Interactors that are involved in protein transport include translocon-related proteins (SSR1, SRP68), a COPII subunit (SEC16A), COPI subunits (COPA, COPG2), and an endocytosis-related protein (CLINT1).

In addition, we identified numerous proteolysis-related interactors. These include ERAD factors, including a number of ubiquitin E3 ligase complexes (HUWE1, TRIM21, UBR5, LTN1, UBE3C, CAND1, KLHL22 and DDB1), an ubiquitin E1 activating enzyme (UBA1), a deubiquitination enzyme (USP9X), retrotranslocation proteins (SEL1L and VCP), and other factors (ERLIN2). As expected, since the α1(A322D) protein undergoes fast ERAD, a number of ERAD factors were recognized in the top left corner in the 2D plot (**Figure 3B**).

### Verification of the role of selected GABA_A_ receptors proteostasis network components

The interactions between selected GABA_A_ receptor proteostasis network components and WT α1 and α1(A322D) variant were verified using co-immunoprecipitation and Western blot analysis. We focus on molecular chaperones and ERAD factors due to their critical role in the protein quality control of GABA_A_ receptors in the ER. Previously, we showed that BiP (HspA5), an Hsp70 family chaperone in the ER lumen, interacts with WT α1 and α1(A322D) subunits and promotes their maturation in the ER (34,43,44). Here, we verified that HspA8, a Hsp70 family chaperone in the cytosol, and its co-chaperones, DNAJA1 and DNAJA2, interact with WT α1 and α1(A322D) subunits (**Figure 4A**), suggesting that Hsp70s and their Hsp40 co-chaperones coordinate the folding of GABA_A_ receptors both in the ER and in the cytosol. Previously, we showed that calnexin (CANX) interacts with WT α1 and α1(A322D) subunits in a glycosylation-dependent manner (34,43,44). Here, we verified that UGGT1, a reglucosylation enzyme that rescues glycoproteins with minor folding defects, interacts with α1 subunits (**Figure 4A**), suggesting its potential role in folding glycosylated GABA_A_ receptors. In addition, the interaction between α1 subunits and UNC45A, an Hsp90 co-chaperone, was demonstrated (**Figure 4A**).

**Figure 4.**
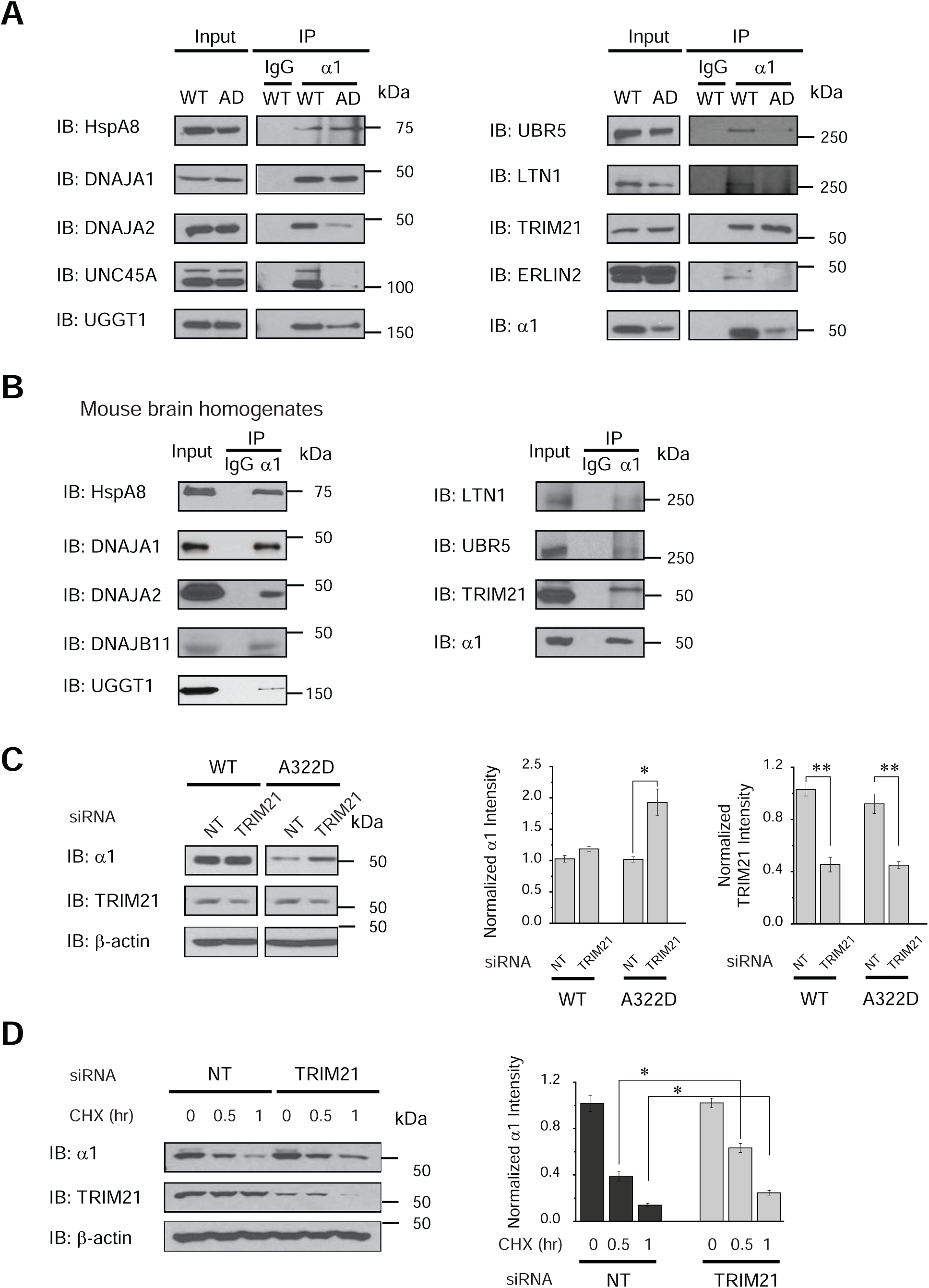
Verification of the role of selected GABA_A_ receptor interactors. HEK293T cells stably expressing WT α1β2γ2 or α1(A322D)β2γ2 GABA_A_ receptors (**A**) or mouse brain homogenates (**B**) were immunoprecipitated using an antibody against α1 subunits and then subjected to SDS-PAGE and Western blot analysis. Three replicates were carried out. AD, α1(A322D) variant; IP, immunoprecipitation; IB, immunoblotting. IgG serves as a negative control during the co-immunoprecipitation. (**C**) HEK293T cells stably expressing WT α1β2γ2 or α1(A322D)β2γ2 GABA_A_ receptors were transfected with non-targeting (NT) siRNA or siRNA against TRIM21. Forty-eight hours post transfection, cells were lysed and subjected to Western blot analysis. Quantification of the normalized protein band intensity was shown on the right (n = 3). β-actin serves as a loading control. (**D**) HEK293T cells expressing α1(A322D)β2γ2 receptors were transfected as in (**C**). Cycloheximide (CHX), a potent protein synthesis inhibitor, was added to the cell culture medium for the indicated time before cell lysis and Western blot analysis. Quantification of the normalized remaining α1 band intensity was shown on the right (n = 3). *, *p* < 0.05; **, *p* < 0.01.

Furthermore, we verified the cellular interactions between GABA_A_ receptors and three ubiquitin E3 ligases that we identified (LTN1, UBR5, and TRIM21) (**Figure 4A**). Mammalian ubiquitin E3 ligases play a central role in the ubiquitination and targeting of their client proteins to the cellular clearance pathway (45-47). Therefore, these E3 ligases are promising candidates to direct GABA_A_ receptors to the proteasome for degradation. In addition, the interaction between α1 subunits and ERLIN2, a potential ERAD factor, which was known to promote the degradation of inositol 1,4,5-trisphosphate (IP_3_) receptors on the ER membrane (48), was confirmed (**Figure 4A**).

To evaluate whether the interactions between GABA_A_ receptors and the proteostasis network components that we identified are utilized in the mammalian central nervous system, we used mouse brain homogenates to carry out co-immunoprecipitation assay. Indeed, we demonstrated the endogenous interactions between α1 subunits and their selected interactors (**Figure 4B**), consistent with the result that most interactors are well conserved between HEK293 cells and the nervous system (**Figure 2C**).

We selected TRIM21, an ubiquitin E3 ligase, for more detailed study. Knocking down TRIM21 using siRNA in HEK293T cells significantly increased the total α1(A322D) protein level (**Figure 4C**, cf. lane 4 to lane 3) without an apparent influence on WT α1 subunits (**Figure 4C**, cf. lane 2 to lane 1), indicating that TRIM21 preferably targets misfolded α1(A322D) subunits for degradation. Moreover, cycloheximide, a potent protein synthesis inhibitor, was applied to the cells for the indicated time to determine the degradation kinetics of α1(A322D) proteins. Compared to non-targeting siRNA control, cycloheximide-chase experiments showed that knocking down TRIM21 increased the remaining α1(A322D) protein levels from 39% to 63% at 0.5 h post cycloheximide application (**Figure 4D**, cf. lane 2 to lane 5), and from 14% to 25% at 1 h post cycloheximide application (**Figure 4D**, cf. lane 3 to lane 6), indicating that depleting TRIM21 attenuated the degradation of α1(A322D) subunits substantially. These results supported that TRIM21 positively regulates the ERAD of the misfolding-prone α1(A322D) subunits.

## Discussion

The comprehensive profiling of the interactomes for GABA_A_ receptors enables us to assemble a cellular model about how the proteostasis network regulates their biogenesis (**Figure 5**). GABA_A_ receptor subunits are co-translationally targeted to the ER membrane (**Figure 5**, State 1), which is mediated by the signal recognition particle (SRP) complex (49). The insertion of the transmembrane domains of GABA_A_ receptors into the lipid bilayer could be facilitated by the SEC61 translocon as well as the ER membrane complex (EMC) (50-52). Our SILAC analysis identified SRP68 and SSR1, which is the α subunit of a translocon-associated protein. GABA_A_ receptor subunits have several Asn *N*-linked glycosylation sites in the ER lumen, which are in the Asn-X-Ser/Thr sequence motif (X can be any residue except Pro): the α1 subunit has two sites at N38 and N138; the β2 subunit has three sites at N32, N104, and N173; and the γ2 subunit has three sites at N52, N129, and N247. The oligosaccharyltransferase (OST) complex is responsible for the transfer of 14-monosaccharide residues Glc3Man9GlcNAc2 (Glc: glucose, Man: mannose, GlcNAc: N-acetylglucosamine) to the above Asn residues in GABA_A_ receptors (**Figure 5**, State 2) (53). Our SILAC analysis identified three OST subunits, including DDOST, RPN1 and RPN2. *N*-linked glycans serve as protein maturation and quality control tag (54). The *N*-linked glycan is then trimmed by α-glucosidase I and α-glucosidase II sequentially to create monoglucosylated glycan (**Figure 5**, State 3). Our SILAC analysis identified GANAB, the α subunit of ER α-glucosidase II. The monoglucosylated GABA_A_ receptor subunit is the client of the membrane-bound lectin chaperone calnexin (CANX) for the folding process (**Figure 5**, State 3 to State 4). In addition, heat shock proteins facilitate the folding of GABA_A_ receptor subunits in a glycan-independent manner both in the ER lumen and in the cytosol (**Figure 5**, State 2 to State 4). Our SILAC analysis identified major Hsp60, Hsp70, Hsp90 family chaperones as well as their co-chaperones, such as HspA5 (BiP) and its co-chaperone DNAJB11 (ERdj3), and HspA8 (Hsc70) and its co-chaperones DNAJA1/2/3. Since each GABA_A_ subunit has a signature Cys-loop in the ER lumen, protein disulfide isomerases (PDIases) play a critical role in assisting the formation of proper disulfide bonds. Our SILAC analysis identified P4HB, an ER PDIase.

**Figure 5.**
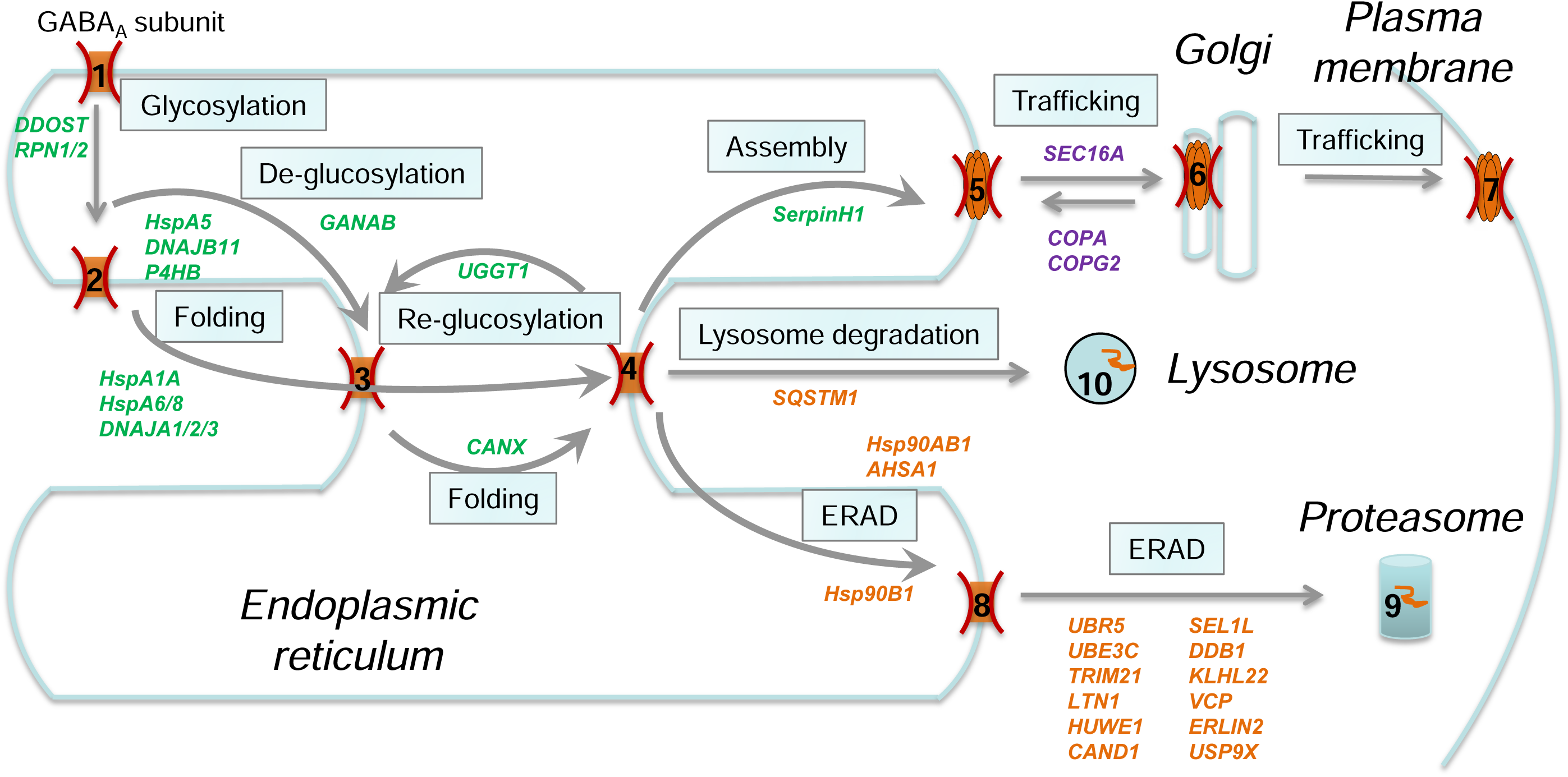
Cellular model of the proteostasis network for GABA_A_ receptors. Interactors that were identified from our SILAC analysis are included in each step. Interactors that are expected to play a role in the folding and assembly of GABA_A_ receptors are colored in green; those in their degradation, in orange; and those in their transport, in purple. The GABA_A_ receptor subunit is co-translationally translocated to the ER membrane (State 1). The OST complex, including DDOST, RPN1, and RPN2, installs *N*-linked glycans to the subunit (State 1 to State 2). The glycoprotein is trimmed by ER α-glucosidases, such as GANAB (State 2 to State 3), and subjected to calnexin (CANX) folding cycles in a glycan-dependent manner (State 3 to State 4). In parallel, heat shock proteins and their co-chaperones facilitate the folding of the subunit both in the ER lumen (HspA5, DNAJB11, P4HB) and in the cytosol (HspA1A, HspA6, HspA8, DNAJA1, DNAJA2, DNAJA3) (State 2 to State 4). After coordinated folding efforts (State 4), the subunit faces three possible routes. *First*, if folded into the native state, the subunit assembles with other subunits to form a heteropentamer with the assistance from assembly factors, such as SerpinH1 (State 4 to State 5). The assembled receptors engage the COPII machinery, including SEC16A, for anterograde trafficking to the Golgi and to the plasma membrane (State 5 to State 6 and State 7). The COPI machinery, including COPA, COPG2, aids the recycle of the subunits from the Golgi back to the ER (State 6 to State 5). *Second*, non-native subunits are reglucosylated by UGGT1 for additional folding attempts (State 4 to State 3). *Third*, terminally misfolded subunits undergo ERAD pathway, being recognized, ubiquitinated, dislocated, and targeted to the proteasome for degradation (State 4 to State 8 and State 9). Ubiquitin E3 ligases (UBR5, UBE3C, TRIM21, LTN1, HUWE1) and other ERAD factors (Hsp90B1, Hsp90AB1, AHSA1, SEL1L, VCP, ERLIN2) potentially play a critical role in this process. Aggregated subunits could utilize the lysosome-related degradation pathway (State 4 to State 10).

After the collaborative folding efforts (**Figure 5**, State 4), GABA_A_ receptor subunits have three possible pathways. *First*, if they achieve the natively folded state, they can assemble with other subunits to form a pentameric receptor for further anterograde trafficking (**Figure 5**, State 4 to State 5). Our SILAC analysis identified SerpinH1, a molecular chaperone that has the potential to facilitate the assembly process (unpublished result). Properly assembled pentameric GABA_A_ receptors engage the trafficking factors to exit the ER, traffic through the Golgi, and travel en route to the plasma membrane (**Figure 5**, State 5 to 6 and 7). Our SILAC analysis identified SEC16A in the COPII machinery that regulates the anterograde cargo protein vesicle transport from the ER to the Golgi (55), as well as COPA and COPG2 in the COPI machinery that regulates the retrograde retrieval of cargo proteins from the Golgi to the ER for recycling (56). *Second*, non-natively folded subunits are recognized by UDP-glucose glycoprotein glucosyltransferase 1 (UGGT1), which was identified from our SILAC analysis. UGGT1 acts as a folding sensor and reglucosylates its substrates for re-entering the calnexin folding cycles for additional folding attempts (**Figure 5**, State 4 to State 3) (57). *Third*, terminally misfolded subunits are subjected to either the ERAD clearance pathway or the lysosome-related degradation pathway (58). During the ERAD, misfolded subunits are recognized, ubiquitinated, dislocated from the ER into the cytosol, and targeted to the proteasome for degradation (**Figure 5**, State 4 to 8 and 9). Our SILAC analysis identified ubiquitin E3 ligases (HUWE1, UBR5, UBE3C, TRIM21, LTN1) and other important ERAD factors, such as SEL1L and VCP. If misfolded subunits tend to form large aggregates in the ER, they are likely to be targeted to the lysosome for degradation (**Figure 5**, State 4 to State 10). Several lysosomal degradation pathways have been described, including macroautophagy, selective ER-phagy, and direct ER-to-lysosome-associated degradation (59). Our SILAC analysis identified SQSTM1 (p62), a marker protein during autophagy by acting as an autophagosome cargo protein.

Previously, we reported that a number of proteostasis network components regulate the folding, degradation and trafficking of GABA_A_ receptors (34,43,60-62), and most of them, such as BiP (HspA5), calnexin (CANX), Grp94 (Hsp90B1), VCP, and SEL1L, were identified from this quantitative SILAC-based IP-MS/MS analysis, indicating the effectiveness of this proteomics approach. For example, we showed that BiP and calnexin interacted with both WT α1 and α1(A322D) subunits; further, overexpression of BiP or calnexin was sufficient to enhance the ER-to-Golgi trafficking efficiency of WT α1 and α1(A322D) subunits (34). Also the interaction between calnexin and α1 subunits is dependent on the *N*-linked glycosylation since mutating the glycosylation sites in α1 subunits substantially decreases such interaction (34,44). These results indicated that BiP and calnexin act as pro-folding chaperones to facilitate the productive folding of GABA_A_ receptors. Regarding the ERAD factors, previously we showed that Grp94 interacted with α1 subunits and that depleting Grp94 decreased ubiquitinated α1 subunits and their degradation rate (60). In addition, we demonstrated that knocking down SEL1L (a critical co-factor of the ubiquitin E3 ligase Hrd1 complex) or VCP (an ATPase that extracts misfolded proteins from the ER membrane to the cytosol) increased the total protein levels of α1(A322D) subunits (43,62). These results supported their critical role in targeting misfolded GABA_A_ receptors to the ERAD pathway. Such knowledge assisted the integration of the interactors into the proteostasis network for GABA_A_ receptors (**Figure 5**).

Additionally, the role of novel interactors of GABA_A_ receptors in regulating their proteostasis remains to be determined in the future. For instance, reticulocalbin 2 (RCN2), a Ca^2+^-binding protein in the ER lumen, is abundant in the central nervous system, and its gene expression is upregulated in patients with idiopathic absence epilepsies (41). Dedicator of cytokinesis protein 7 (DOCK7), which is localized to the developing axons, activates Rac1 and Rac3 small GTPases and regulates neuronal polarity; moreover, its variants cause developmental and epileptic encephalopathy 23 (DEE23) (63,64). SLC39A7 (ZIP-7) is a Zn^2+^ efflux transporter in the ER membrane; ZIP7 depletion causes neurodevelopmental impairments and ZIP7 signaling pathway provides neuroprotective effects in recurrent seizures (65,66).

Loss of function of GABA_A_ receptors is a prominent cause of genetic epilepsies, and recent advances in genetics have identified an increasing number of GABA_A_ variants that are associated with epilepsy (16,19). Current treatments for epilepsy focus on relieving symptoms instead of targeting the underlying causes, leaving behind much unmet medical needs. Recently, we demonstrated that adapting the ER proteostasis network is a promising strategy to restore the functional surface expression of pathogenic GABA_A_ variants (43,67). Therefore, identification of the proteostasis network for GABA_A_ receptors paves the foundation to adjust the cellular folding, assembly, degradation, and trafficking pathways to fine-tune the functional surface expression of GABA_A_ receptors as a novel therapeutic strategy to ameliorate epilepsy.

### Experimental procedures

#### Plasmids

The pCMV6 plasmids containing human GABA_A_ receptor α1 subunit (Uniprot #: P14867-1) (# RC205390), β2 subunit (isoform 2, Uniprot #: P47870-1) (#RC216424), γ2 subunit (isoform 2, Uniprot #: P18507-2) (#RC209260), and pCMV6 Entry Vector plasmid (pCMV6-EV) (#PS100001) were purchased from Origene. The missense mutation A322D in the GABA_A_ receptor α1 subunit was constructed using a QuikChange II site-directed mutagenesis Kit (Agilent Genomics, #200523), and the cDNA sequences were confirmed by DNA sequencing.

#### Antibodies

The mouse monoclonal anti-GABRA1 antibody (#MAB339) was obtained from Millipore. The rabbit polyclonal anti-DNAJB11 (#GTX105619) antibody was obtained from GeneTex. The rabbit polyclonal anti-ERLIN2 antibody (#PA5-21736) was obtained from Thermo Fisher Scientific. The rabbit polyclonal anti-LTN1 antibody (#AP53685PU-N) was obtained from Novus Biologicals. The rabbit polyclonal anti-DNAJA1 antibody (#AP5849C) and anti-HspA8 (#AP2872A) antibody were obtained from Abgent. The rabbit polyclonal anti-DNAJA2 antibody (#12236-1-AP), anti-TRIM21 antibody (#10108-1-AP), and anti-UNC45A (#15479-1-AP) antibody were obtained from Proteintech. The rabbit monoclonal anti-UGGT1 antibody (#3543-1) was obtained from Epitomics. The mouse monoclonal anti-β-actin antibody (#A1978) was obtained from Sigma. The rabbit polyclonal anti-UBR5 antibody (#A300-573A) was obtained from Bethyl Laboratories.

#### Cell Culture and Transfection

HEK293T cells (ATCC, # CRL-3216) were maintained in Dulbecco’s Modified Eagle Medium (DMEM) (Fisher, #SH3024301) with 10% heat-inactivated fetal bovine serum (Fisher, #SH3039603HI) and 1% Penicillin Streptomycin (Fisher, #SV30010) at 37°C in 5% CO_2_. Monolayers were passaged upon reaching confluency with 0.05% Trypsin protease (Fisher, #SH30236.01). HEK293T cells were grown in 6-well plates or 10-cm dishes and allowed to reach ∼70% confluency before transient transfection using TransIT-2020 (Mirus, #MIR 5400), or siRNA treatment (50 nM) using the HiPerfect Transfection Reagent (Qiagen, #301707), according to the manufacturer’s instruction. The small interfering RNA (siRNA) duplexes were obtained from Dharmacon: TRIM21 (J-006563-10-0005) and Non-Targeting siRNA (D-001810-01-20) as negative control. Forty-eight hours post-transfection, cells were harvested for further analysis.

#### SDS-PAGE and Western Blot

Cells were harvested with 0.05% Trypsin protease (Fisher, #SH30236.01) and then lysed with the lysis buffer (50 mM Tris, pH 7.5, 150 mM NaCl, and 1% Triton X-100) supplemented with complete protease inhibitor cocktail (Roche, #4693159001). Cell lysates were cleared by centrifugation (15,000 × *g*, 10 min, 4^°^C) and the supernatants were collected as total proteins. Protein concentration was determined by MicroBCA assay (Pierce, #23235). Equal amounts of total proteins were separated in 8% reducing SDS-PAGE gels, and Western blot analysis was performed using appropriate antibodies.

#### Mouse brain homogenization

C57BL/6J mice (Jackson laboratory) at 6-10 weeks were sacrificed and the cortex was isolated and homogenized in the homogenization buffer (25 mM Tris, pH 7.6, 150 mM NaCl, 1 mM EDTA, and 2% Triton X-100) supplemented with the Roche complete protease inhibitor cocktail. The sample was centrifuged at 800 × *g* for 10 min at 4^°^C. The pellet was re-homogenized in the same homogenization buffer and centrifuged at 800 × *g* for 10 min at 4^°^C. The combined supernatants were placed on a rotating device for 2 h at 4^°^C and then centrifuged at 15,000 × *g* for 30 min at 4^°^C. The resulting supernatant was collected as mouse brain homogenate, and its protein concentration was determined by a MicroBCA assay (Pierce, #23235). This animal study followed the guidelines of the Institutional Animal Care and Use Committees (IACUC) at Case Western Reserve University, and was carried out in agreement with the recommendation of the American Veterinary Medical Association Panel on Euthanasia.

#### Immunoprecipitation

For immunoprecipitation using cell lysates (500 µg) and the mouse brain homogenates (1 mg), they were pre-cleared with 30 µL of protein A/G plus-agarose beads (Santa Cruz, #sc-2003) and 1.0 µg of normal mouse IgG (Santa Cruz, #sc-2025) for 1 hour at 4°C to remove nonspecific binding proteins. The pre-cleared cell lysates were incubated with 2.0 µg of mouse anti-α1 antibody for 1 hour at 4°C and then with 30 µL of protein A/G plus agarose beads overnight at 4°C. Afterwards, the beads were collected by centrifugation at 8000 × *g* for 30 s, and washed three times with lysis buffer. The complex was eluted by incubation with 40 µL of 2x Laemmli sample buffer (Biorad, #1610737) in the presence of DTT. The immunopurified eluents were separated in 8% SDS-PAGE gel, and Western blot analysis was performed using appropriate antibodies.

#### Stable Isotope Labeling with Amino Acids in Cell Culture (SILAC)-based Quantitative Proteomics Analysis

##### SILAC Labeling

SILAC is an *in vivo* labeling strategy for mass spectrometry-based quantitative proteomics (32,68). HEK293T cells stably expressing either WT α1β2γ2 or α1(A322D)β2γ2 GABA_A_ receptors were labeled with heavy media (SILAC DMEM media (Pierce, #88420) plus 10% dialyzed FBS (Sigma, #F0392), 1% Penicillin Streptomycin, 0.01% ^13^C_6_ L-Lysine-2HCL (Pierce, #89988), 0.01% ^13^C_6_ L-Arginine-HCL (Pierce, #88210), and 0.002% L-Proline (Pierce, #88430)), whereas HEK293T cells that were transfected with empty vector (EV) plasmids were cultured in normal light media (SILAC DMEM media plus 10% dialyzed FBS, 1% Penicillin Streptomycin, 0.01% L-Lysine-2HCL (Pierce, #88429), 0.01% L-Arginine-HCL (Pierce, #88427) and 0.002% L-Proline) for 14 days to ensure complete labeling.

##### Co-immunoprecipitation and In-Gel Digestion

Cells were then harvested with 0.05% Trypsin and lysed in the lysis buffer (50 mM Tris, pH 7.5, 150 mM NaCl, and 1% Triton X-100) supplemented with Roche complete protease inhibitor cocktail. Lysates were cleared by centrifugation (15,000 × *g*, 10 min, 4^°^C). Protein concentration was determined by MicroBCA assay (Pierce). The same amount of light and heavy cell lysates was mixed by 1:1. Six mg of total proteins were immunoprecipitated using a mouse monoclonal anti-GABA_A_ receptor α1 subunit antibody. The immunoisolated complexes were separated by reducing SDS-PAGE and stained with Coomassie blue to visualize protein gel bands (**Figure 1C**). The gel was washed in distilled water to remove excess background stain. The gel was then divided to six parts evenly, excised and destained with 500 μL of 1:1 ACN (acetonitrile) and 100 mM ABC (ammonium bicarbonate) solution for 2-8 hours. Afterwards, 10 mM reductive TCEP (tris(2-carboxyethyl)phosphine) was added for 30 min, and then free cysteines were alkylated with 55 mM IAA (iodoacetamide) for 20 min in the dark. ACN and 100 mM ABC were used to dehydrate and rehydrate the gel pieces alternatively for three times. Gel pieces were swelled in 50 mM ABC containing freshly prepared 10 ng/μL trypsin (Promega, sequencing grade) and digested overnight. Peptides were extracted with 60% ACN/5% FA (formic acid). Digested peptides were cleaned through C18 spin columns (Thermo Pierce, Rockford, IL), dried in SpeedVac, and saved at -80°C if not immediately analyzed.

##### Tandem Mass Spectrometry Analysis

Three biological replicates were analyzed by liquid chromatography-tandem mass spectrometry (LC-MS/MS). Digested peptides were reconstituted with 0.1% formic acid, and analyzed by LC-MS/MS. Separation of peptides via capillary liquid chromatography was performed using Waters nanoAquity system (Waters Corp., Milford, MA). Mobile phase A (aqueous) contained 0.1% formic acid in 5% acetonitrile, and mobile phase B (organic) contained 0.1% formic acid in 85% acetonitrile. Separation was achieved using a C18 column (75 μm × 20 cm, Waters Corp., Ethylene Bridged Hybrid column # BEH300) through a 300-minute gradient of 6% to 45% mobile phase B at a flow rate of 300 nL/min. MS analysis was performed using a hybrid linear ion trap Orbitrap Velos mass spectrometer (LTQ-Orbitrap Velos, Thermo, Waltham, MA). Survey scans were operated at 60,000 resolution, followed by twenty CID (collision-induced dissociation) fragmentations.

##### Database Search

Acquired tandem mass spectra were searched against Uniprot human protein database with 20,244 protein entries, downloaded on 5/5/2011. A decoy database containing the reversed sequences of all proteins was appended to estimate the false discovery rate (69). Protein identification using Sequest (70) or ProLuCID (71) and DTASelect (72,73) and quantification using Census (74) were performed using the Integrated Proteomics Pipeline (IP2, Integrated Proteomics Applications, San Diego, CA). Mass accuracy was limited to 10 ppm for precursor ions and 0.6 Da for product ions, with tryptic enzyme specificity and up to two missed cleavages. Static modifications included carbamidomethylation on cysteines (57 Da), and variable modifications included oxidation on methionines (16 Da) in addition to the SILAC modifications, ie., heavy ^13^C_6_ on lysines and arginines (6 Da). DTASelect (72,73) was applied to generate search results for peptide-to-spectra matches (PSMs) with a maximum false discovery rate (FDR) of 5%, yielding a protein FDR of less than 1%. Protein quantification based on SILAC peptide pairs was performed with Census (74). Census allows users to filter peptide ratio measurements based on a correlation threshold because the correlation coefficient (values between zero and one) represents the quality of the correlation between the unlabeled and labeled chromatograms and can be used to filter out poor quality measurements. In this study, only peptide ratios with correlation values greater than 0.5 were retained. High-confidence protein quantifications were based on two or more SILAC pairs for a given protein.

The interactomes of α1 subunits and α1(A322D) subunits of GABA_A_ receptors were identified using the SILAC ratio with arbitrary yet strict criteria to remove potential false positives. Only proteins with *p* < 0.05 were further considered. To be included as an interactor, it must (1) have a SILAC ratio of WT α1/EV or α1(A322D)/EV to be at least 1.30; and (2) have a Benjamini and Hochberg correction (35) of false discovery rate (FDR) of no more than 0.10. For display purpose in **Figure 3B**, if the SILAC ratio is greater than 4.0, it is displayed as 4.0; if a protein is only detected in one sample, the SILAC ratio of this protein in the other sample is artificially set to 1.0. The final list of the interactomes of α1 subunits and α1(A322D) subunits of GABA_A_ receptors is presented in Supplementary **Table S1**.

#### Gene Ontology Analysis

Cellular Component and Biological Process of GABA_A_ receptor interactomes were analyzed using Database for Annotation, Visualization and Integrated Discovery (DAVID) (https://david.ncifcrf.gov/) (39,40).

Protein abundance analysis was performed using the ProteomicsDB database (https://www.proteomicsdb.org/) (36). RNA abundance analysis was carried out using the Human Protein Atlas database (https://www.proteinatlas.org/) (37,38).

### Cycloheximide (CHX)-chase assay

HEK293T cells stably expressing α1(A322D)β2γ2 GABA_A_ receptors were seeded at 2.5 × 10^5^ cells per well in 6-well plates and incubated at 37°C overnight. Cells were then transfected with indicated siRNA. Forty-eight hours post transfection, cells were incubated with cycloheximide (100 μg/mL) (Enzo, #ALX-380-269-G001), a potent protein synthesis inhibitor, to stop protein translation, and chased for the indicated time. Cells were then harvested, and lysed for SDS-PAGE and Western blot analysis.

### Statistical analysis

All data are presented as mean ± SEM. Statistical significance was calculated using two-tailed Student’s t-Test for two group comparison. If more than two groups were compared, one-way ANOVA followed by post-hoc Tukey test was used. A *p* value of less than 0.05 was considered statistically significant. *, *p* < 0.05; **, *p* < 0.01.

### Data availability

All data are contained within the manuscript.

## Supporting information

This article contains supporting information. Supporting information includes six supplemental tables.

## Acknowledgements

This work was supported by the National Institutes of Health (R01NS105789 and R01NS117176 to TM) and Clinical Translational Science Collaborative of Cleveland CTSA (UL1RR024989) from the National Center for Research Resources and the National Center for Advancing Translational Sciences of the NIH.

## Author contributions

Conceptualization, YW and TM; Data curation: YW and XD; Formal analysis: YW and TM; Funding acquisition: TM; Supervision: YW and TM; Writing – original draft: YW and TM; Writing – review & editing: YW, XD, and TM.

## Funding and additional information

The content is solely the responsibility of the authors and does not necessarily represent the official views of the National Institutes of Health.

## Supplemental Tables

**Table S1**. List of proteins and their SILAC ratios from the quantitative proteomics experiments to identify GABA_A_ receptor interactomes. Labeled with a unique entry number for each protein, first 54 proteins overlap within WT α1 interactome and the α1(A322D) interactome. Number 55 to 125 are WT α1 only interactors, whereas number 126 to 176 are α1(A322D) only interactors.

**Table S2**. List of protein abundance and RNA abundance of GABA_A_ receptor interactors. Scores of protein abundance were retrieved from the ProteomicsDB database (https://www.proteomicsdb.org/). Protein comparison included data from HEK293 cells and human brain tissues. Scores of RNA abundance were retrieved from the Human Protein Atlas database (https://www.proteinatlas.org/). RNA comparison included data from HEK cells, SH-SY5Y cells and human brain tissues. Fpkm, fragments per kilobase of exon per million mapped fragments.

**Table S3**. List of cellular location and membrane proteins of GABA_A_ receptor interactors. Entry number for each protein was the same as in **Table S1** for overlapping, WT α1 only, or α1(A322D) only interactors. In the column of *Membrane protein, 1* means that the corresponding protein is indeed an integral membrane protein, whereas *0* means otherwise. Moreover, cellular location statistics and membrane protein statistics were presented as well, summarizing number of interactors in each GO term.

**Table S4**. List of biological process for GABA_A_ receptor interactors. Entry number for each protein was the same as in **Table S1** for overlapping, WT α1 only, or α1(A322D) only interactors. In the column of *Function 2*, further available information was provided. Moreover, biological process statistics were presented as well, summarizing number of interactors in each category.

**Table S5**. List of enriched biological process for GABA_A_ receptor interactors. Data were analyzed and retrieved from Database for Annotation, Visualization and Integrated Discovery (DAVID). Five functional clusters were presented from both WT α1 and α1(A322D) interactors. GO terms, enriched genes, and FDR were included.

**Table S6**. List of the proteostasis network components among GABA_A_ receptor interactors. Based on GO analysis and literature mining, three categories were included as the following: protein folding, proteolysis, and protein transport, for overlapping, WT α1 only, or α1(A322D) only interactors.

